# Quantitative proteome-based guidelines for intrinsic disorder characterization

**DOI:** 10.1101/032847

**Authors:** Michael Vincent, Mark Whidden, Santiago Schnell

## Abstract

Intrinsically disordered proteins fail to adopt a stable three-dimensional structure under physiological conditions. It is now understood that many disordered proteins are not dysfunctional, but instead engage in numerous cellular processes, including signaling and regulation. Disorder characterization from amino acid sequence relies on computational disorder prediction algorithms. While numerous large-scale investigations of disorder have been performed using these algorithms, and have offered valuable insight regarding the prevalence of protein disorder in many organisms, critical proteome-based descriptive statistical guidelines that would enable the objective assessment of intrinsic disorder in a protein of interest remain to be established. Here we present a quantitative characterization of numerous disorder features using a rigorous non-parametric statistical approach, providing expected values and percentile cutoffs for each feature in ten eukaryotic proteomes. Our estimates utilize multiple *ab initio* disorder prediction algorithms grounded on physicochemical principles. Furthermore, we present novel threshold values, specific to both the prediction algorithms and the proteomes, defining the longest primary sequence length in which the significance of a continuous disordered region can be evaluated on the basis of length alone. The guidelines presented here are intended to improve the interpretation of disorder content and continuous disorder predictions from the proteomic point of view.

## 1. Introduction

Once translated, many nascent unfolded polypeptides fold into a highly ordered conformation. However, within the last two decades it has been become increasingly apparent that not all proteins fold into a stable globular structure [1-3]. Rather, many proteins and/or protein regions are thought exhibit intrinsic disorder. Intrinsically disordered proteins or protein regions are those that lack a stable three-dimensional structure under physiological conditions, but instead, exist in a natively unfolded state. From a physicochemical standpoint, disordered regions are often characterized by low complexity and the absence of secondary structure, and often consist of residues with low hydrophobicity and high polarity and charge [4]. Disorder has emerged as a prevalent and important feature in the proteomes of many prokaryotes and eukaryotes. Regarding the latter, it has been estimated that 15-45% of eukaryotic proteins contain “significant” long disordered regions, commonly defined as a disordered stretch of 30 or more amino acids in length [5].

While writing off intrinsically disordered proteins as lacking function would be easy due to the absence of a well-defined tertiary structure, a growing body of evidence supports intrinsically disordered proteins playing important functional roles in various signaling and regulatory processes [4, 6, 7], including apoptosis [8, 9], and cell cycle regulation [10]. Interestingly, disorder may also serve as a recognizable feature. Ube2W, a unique ubiquitin-conjugating enzyme (E2) that mono-ubiquitinates the amino-terminus of target substrates, was recently found to specifically recognize substrates with disordered N-termini *in vitro* [11]. Additional support has been established *in vivo* in a Ube2W knockout mouse model, where both full-length and N-terminal disorder were found to be more prevalent in a subset of testicular proteins exhibiting a 1.5X expression increase in the knock-out compared to wild-type [12]. Some proteins involved in protein misfolding diseases are now understood as being intrinsically disordered as well, including the Amyloid-β peptide in Alzheimer’s disease and α-synuclein in Parkinson’s disease [13].

While analyzing the role of disorder within a single protein or a small set of related proteins is important for understanding the contributions of disorder to protein structure (or the lack of structure) and function, studies must be carried out at the proteomic level to establish critical reference points for disorder characterization. Indeed, proteomic investigations of disorder have been performed and have offered valuable insight into the prevalence of disorder in many organisms [14-16]. However, these studies have not provided guidelines in the form of explicit descriptive statistics, specific to both proteomes and disorder prediction tools, for identifying anomalous disorder features with respect to whole proteomic populations. Without these guidelines in hand, it remains very difficult to understand whether or not a given disorder measure is significant with respect to the population. Guidelines of this nature would be analogous to clinical guidelines used to identify and evaluate whether an individual is overweight or obese based on the body mass index distribution in the population [17-19]. For example, if a protein of interest is found to contain a disordered region that is 25 amino acids in length, is this significant? And how does the context of the primary sequence length influence the evaluation of significance? Before these questions can be answered objectively, a rigorous descriptive statistical analysis of disorder content and continuous disorder must be conducted at the proteome level.

Motivated by these considerations, we analyzed disorder in the proteomes of ten eukaryotic model organisms using a non-parametric descriptive statistical approach. Disorder was estimated using two reputable disorder prediction algorithms, IUPred and DisEMBL, which have a physicochemical basis. While larger-scale disorder studies have been performed, limiting our study to a manageable number of common eukaryotes allowed us to ascertain the quality of the protein sequence pool, quantitatively and qualitatively inspect the accuracy of our statistical methodology, and present objective guidelines for disorder classification in an explicit fashion. This work provides one of the most systematic non-parametric efforts toward standardizing disorder content and continuous length disorder that has been described in the literature.

## 2. Materials and Methods

### 2.1 Proteomes and protein sequences

Primary sequences for all proteins included in our analysis were obtained from UniProt reference proteome files [20]. The ability to visualize data distributions in our study is extremely important for testing and presenting the validity of our nonparametric statistical approach, thereby limiting our study to the proteomes of ten model eukaryotes. Specifically, the *Saccharomyces cerevisiae, Dictyostelium discoideum, Chlamydonmonas reinhardtii, Drosophila melanogaster, Caenorhabditis elegans, Arabidopsis thaliana, Danio rerio, Mus musculus, Homo sapiens, and Zea mays* proteomes were included in our investigation (proteome presentation order was decided by final protein population size, which is described in detail in Section 2.2.). In an effort to obtain the most accurate results possible, only proteins with completely defined primary sequences were considered eligible for our analysis. Proteins with undetermined/unknown, ambiguous, and/or unique amino acids (B, J, O, U, X, Z) were excluded on the basis that the handling of these residues varies greatly among disorder prediction algorithms. A summary of the eligible and ineligible protein populations is displayed in **Table 1**. For a complete list of UniProt accession numbers for all eligible and ineligible proteins, please refer to **Supplemental Table 1**.

**Table 1.** Eligibility screening summary of proteins in each studied proteome.

| Organism | Initial Total | Eligible | Ineligible |
| --- | --- | --- | --- |
| *Saccharomyces cerevisiae* | 6,721 | 6,721 (100%) | 0 (0%) |
| *Dictyostelium discoideum* | 12,746 | 12,733 (99.82%) | 13 (0.18%) |
| *Chlamydomonas reinhardtii* | 14,337 | 14,319 (99.87%) | 18 (0.13%) |
| *Drosophila melanogaster* | 22,024 | 21,673 (98.40%) | 351 (1.60%) |
| *Caenorhabditis elegans* | 26,163 | 26,161 (99.99%) | 2 (0.01%) |
| *Arabidopsis thaliana* | 31,551 | 31,548 (99.99%) | 3 (0.01%) |
| *Danio rerio* | 41,001 | 38,192 (93.15%) | 2,809 (6.85%) |
| *Mus musculus* | 45,263 | 42,306 (93.47%) | 2,957 (6.53%) |
| *Homo sapiens* | 68,485 | 61,423 (89.69%) | 7,062 (10.31%) |
| *Zea mays* | 58,493 | 58,455 (99.94%) | 38 (0.06%) |

Primary sequences were obtained from UniProt reference proteome files. Proteins with undetermined, ambiguous, and/or rare amino acid residues were excluded from our analysis. Initial total, included, and excluded protein sequence counts are displayed for each organism, as well as the percentages of the initial total that have been included and excluded.

### 2.2 Sequence redundancy and uncertainty reduction

While the aforementioned eligibility screening procedure filtered out sequences that are incompatible for disorder prediction, redundant sequences and sequences that are uncertain to exist still remained in the eligible sequence population for each proteome. In order to minimize redundancy and uncertainty within the population of eligible sequences we conducted the following two-step procedure. First, UniRef100 reference cluster information was obtained via the UniProt identification mapping service (accessed programmatically on January 6, 2016) and was used to remove redundant sequences from each proteome [20, 21]. The resulting proteome populations were comprised of (i) UniProt accession numbers of eligible proteins that correspond to the unique set of UniRef100 records found to map *directly* to the reference proteome file, and (ii) UniProt accession numbers of eligible proteins that were found to map to a UniRef100 record that was not contained within the specific reference proteome file. Second, proteins with a UniProt protein existence qualifier of five were subsequently removed, as the existence of these proteins is uncertain [20]. The final population sizes have been displayed for each proteome in **Table 2** (the population size of each proteome was used to determine presentation order, with *Zea mays* representing the largest population in our study following the reduction procedure). The UniProt accession numbers comprising the final population have been included in **Supplemental Table 2** along with the UniProt accession numbers of proteins that have been excluded on the basis of existence uncertainty.

**Table 2.** Summary of redundancy and uncertainty reduction in the number of proteins in each proteome.

| Organism | RR | PE 5 | Final |
| --- | --- | --- | --- |
| *Saccharomyces cerevisiae* | 6,628 | 766 | 5,862 |
| *Dictyostelium discoideum* | 12,688 | 22 | 12,666 |
| *Chlamydomonas reinhardtii* | 14,229 | 1 | 14,228 |
| *Drosophila melanogaster* | 19,640 | 0 | 19,640 |
| *Caenorhabditis elegans* | 23,369 | 7 | 23,362 |
| *Arabidopsis thaliana* | 30,814 | 112 | 30,702 |
| *Danio rerio* | 34,854 | 0 | 34,854 |
| *Mus musculus* | 37,192 | 7 | 37,185 |
| *Homo sapiens* | 51,701 | 588 | 51,113 |
| *Zea mays* | 54,133 | 1 | 54,132 |

### 2.3 Disorder prediction and analysis

Residue-specific disorder scores were obtained using the IUPred long [22, 23] and DisEMBL [24] *ab initio* disorder prediction algorithms. IUPred predicts disorder by estimating the capacity of a protein to form stabilizing interresidue contacts [22, 23]. DisEMBL predicts disorder using three prediction methods based on artificial neural networks – Coils (DisEMBL-C), Hotloops (DisEMBL-H), and Rem465 (DisEMBL-R). DisEMBL-C uses secondary structure prediction to assign residues as belonging to coils if the residues do not fall in α-helix, 310-helix, or β-strand secondary structures [24]. Furthermore, DisEMBL-H predicts disorder by analyzing the subset of DisEMBL-C predictions that have high α-carbon temperature factors, whereas DisEMBL-R has been trained on missing X-ray crystallography coordinates in the Protein Data Bank [24]. Due to DisEMBL-C predictions being an overestimate of disorder (as described by [24]), only results from the DisEMBL-H and DisEMBL-R methods were analyzed for DisEMBL (however, DisEMBL-C probability densities have been included in **Supplemental Fig. 1, 3**, and **4**). Each residue was classified as either “ordered” or “disordered” using algorithm-specific default threshold values [22-24]. Disorder was characterized in each proteome by assessing the disorder content and continuous disorder (CD) distributions. Percent disorder was calculated as the percentage of disordered residues in a protein divided by the protein length, multiplied by one hundred. A CD segment was defined as any stretch of two or more consecutive amino acids having disorder scores above the algorithm-specific threshold value. The minimum length of a CD segment of two amino acids is purely theoretical and we acknowledge that a CD segment of this length might not be structurally important. However, this theoretical minimum was utilized in order to include all possible lengths of predicted CD segments and avoid imposing a subjective minimum length that, if inaccurate, could lead to the exclusion of valid short CD segments from our statistical analysis.

### 2.4 Statistical methods

Due to the lack of normality in many of the distributions examined (**Supplemental Fig. 1, 3**, and **4**), we utilized a non-parametric statistical approach. Kernel density estimation (KDE) with renormalization was used to approximate the probability density function (PDF); the PDF approximation is based on the method of Jones [25]. For distributions of percentages, the PDF was approximated on the bounded domain of zero to one hundred, inclusive. For non-percentage data, a bounded domain defined by the minimum and maximum values was used to approximate the PDF.

To determine the expected value (*E(x)*), the PDF (*f(x)*) was integrated via Eq. 1 (‘lb’ and ‘ub’ represent the lower and upper bound, respectively):

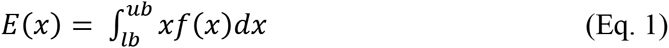

The interquartile range (bounded by the 25^th^ and 75^th^ percentiles) was used to examine the dispersion of the data. In general we interpret disorder content below the 25^th^ percentile to be significantly ordered, with features above the 75^th^ percentile to be significantly disordered. While this percentile-based approach may appear naïve given that it assumes equal proportions of proteins with significantly ordered and disordered features in each proteome, assessing the 25^th^ and 75^th^ percentiles allows for the identification of values where a feature begins to depart the central 50% of the population and therefore provides a descriptive statistical guideline for interpreting the output of specific computational disorder prediction tools. However, this approach cannot be used to interpret order in IUPred disorder content distributions, due to the extremely low 25^th^ percentile values observed. In order to provide cutoffs for gauging extreme order and disorder, the 5^th^ and 95^th^ percentile values have been reported as well.

### 2.5 Computational analysis

All internal noncommercial software created for use in our investigation was written in Python 2.7.10. Results were stored in a SQLite3 database. The database is available upon request.

## 3. Results

### 3.1 Disorder content varies approximately from ∼14-39% depending on predictor, with the majority of the examined proteomes having expected values below 30%

In order to obtain an overall summary of disorder, we first assessed the distribution of disorder percentages in each proteome. To accomplish this, disorder scores were obtained for all residues using each of the aforementioned disorder prediction algorithms. Percent disorder was calculated as described in the **Materials and Methods**. Kernel density estimation was used to approximate the probability density function, which was then integrated using **Eq. 1** to obtain the expected value.

The disorder content distribution is positively skewed in each of the ten eukaryotes analyzed. Expected values were found to range between ∼17-27%, ∼30-39%, and ∼14-22% for IUPred, DisEMBL-H, and DisEMBL-R, respectively (**Fig. 1**). Whereas the human disorder content distribution exhibited the greatest dispersion for DisEMBL predictions (**Fig. 1B, C**), the IUPred distributions did not follow this trend, as the greatest spread was found in the *Drosophila melanogaster* proteome (**Fig. 1A**). Nevertheless, the IUPred and DisEMBL-R predictions were consistent with the 20.5% average disorder percentage recently reported by an investigation of disorder in 110 eukaryotes using IUPred and Espritz [16], as well as an earlier proteomic study conducted using DISOPRED2 that found disorder content to vary from ∼16-22% in five eukaryotes [14]. DisEMBL-H predictions were found to be much higher overall, but still in agreement with the 35-45% range reported by [15], which utilized the PONDR VSL2B predictor. Probability densities are displayed in **Supplemental Fig. 1**. Explicit statistical values have been displayed in **Supplemental Table 3**.

**Fig. 1.**
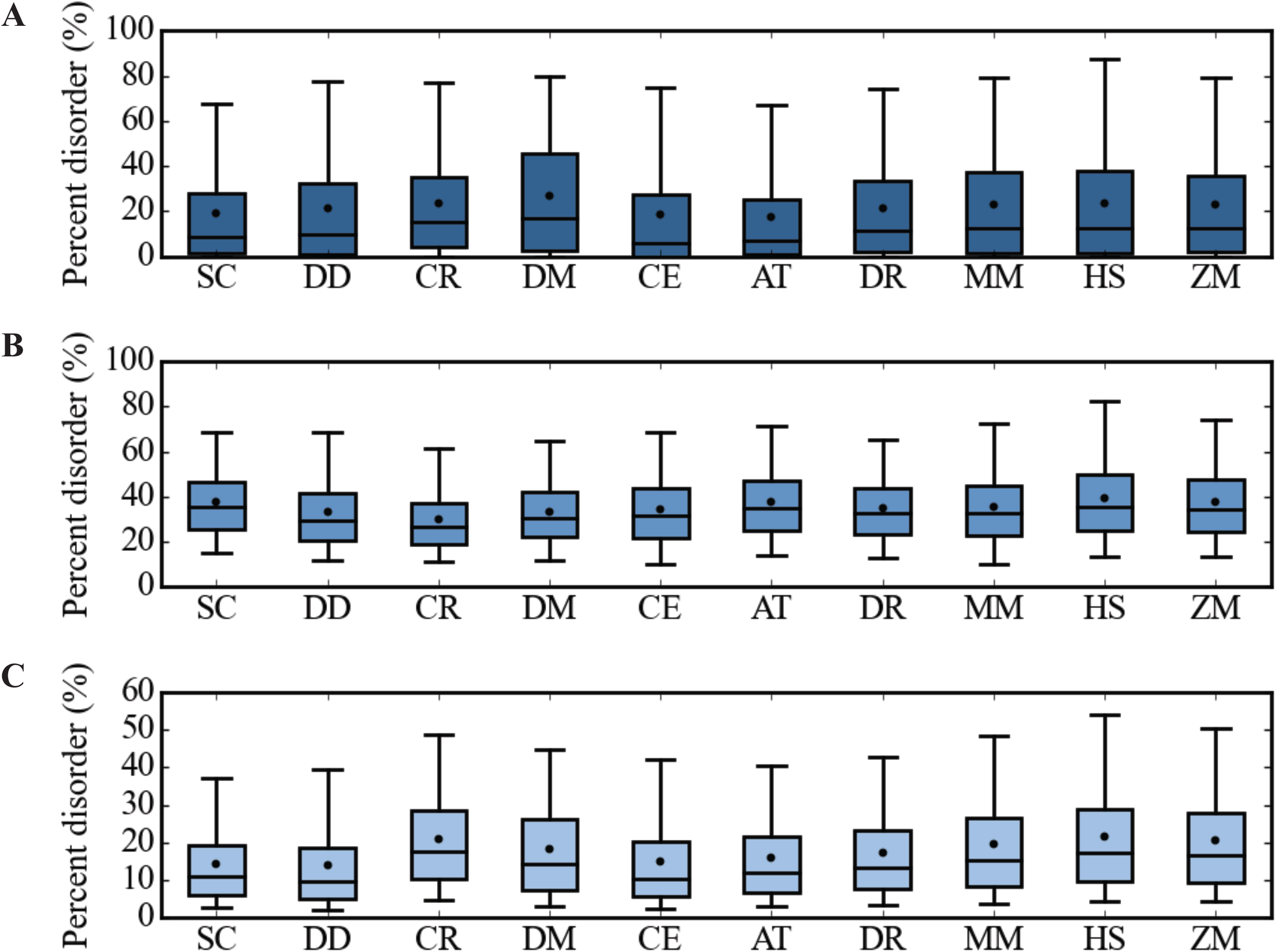
Percent disorder distribution in ten eukaryotic proteomes. Boxplots of percent disorder determined by (A) IUPred, (B) DisEMBL-H, and (C) DisEMBL-R are shown. The horizontal line indicates the median, whereas the dots indicate the expected value determined via **Eq. 1**. The whiskers represent the 5^th^ and 95^th^ percentile values. Numerical values are summarized in **Table 4**.

### 3.2 The majority of the studied eukaryotes contain a least one continuous disorder domain

By definition, a CD region must contain a minimum of two consecutive disordered amino acids. With a length of two amino acids representing the theoretical minimum length of a CD region, completely ordered proteins (0% disorder) and proteins with disorder composition consisting entirely of isolated disordered amino acids must be excluded from our CD analysis. For all eukaryotic proteomes analyzed here, over 70% contain at least one CD stretch as determined by any of the individual prediction methods (**Table 3**). Moreover, disorder percentages within the CD-containing protein populations were nearly identical to those of the entire populations for DisEMBL-H and DisEMBL-R predictions (compare **Supplemental Fig. 2A** to **Fig. 1**), whereas differences were observed for IUPred-predicted disorder with disorder content found to be greater in the CD-containing population (**Supplemental Fig. 2A, B**). This result can be explained by the fact that IUPred predicted a greater amount of isolated disordered amino acids than did DisEMBL, causing the size of the total and CD-containing protein populations to be substantially different for IUPred and identical for DisEMBL (**Table 3**). Nevertheless, provided over two thirds of each proteome exhibited continuous disorder, we reasoned that all of the eukaryotes included in our investigation contained a population of eligible (CD-containing) proteins large enough to determine representative expected values for various CD features.

**Table 3.** Summary of continuous disorder prevalence in ten eukaryotic proteomes.

| Organism | IUPred | DisEMBL-H | DisEMBL-R |
| --- | --- | --- | --- |
| *Saccharomyces cerevisiae* | 4,762 (81.24%) | 5,862 (100.0%) | 5,859 (99.95%) |
| *Dictyostelium discoideum* | 9,806 (77.42%) | 12,661 (99.96%) | 12,643 (99.82%) |
| *Chlamydomonas reinhardtii* | 12,112 (85.13%) | 14,226 (99.99%) | 14,223 (99.96%) |
| *Drosophila melanogaster* | 16,486 (83.94%) | 19,634 (99.97%) | 19,637 (99.98%) |
| *Caenorhabditis elegans* | 16,437 (70.36%) | 23,353 (99.96%) | 23,324 (99.84%) |
| *Arabidopsis thaliana* | 23,939 (77.97%) | 30,698 (99.99%) | 30,693 (99.97%) |
| *Danio rerio* | 28,922 (82.98%) | 34,850 (99.99%) | 34,841 (99.96%) |
| *Mus musculus* | 29,393 (79.05%) | 37,175 (99.97%) | 37,178 (99.98%) |
| *Homo sapiens* | 39,369 (77.02%) | 51,104 (99.98%) | 51,097 (99.97%) |
| *Zea mays* | 43,967 (81.22%) | 54,119 (99.98%) | 54,106 (99.95%) |

### 3.3 For many of the eukaryotes examined, a CD region greater than 30 amino acids is expected, and the length of significantly long disordered stretches varies substantially between predictors

Isolated disordered amino acids and continuous clusters of disordered residues constitute the two most basic disorder arrangements. While isolated disordered residues may influence the structure of some proteins, longer continuously disordered segments provide better indicators of protein regions that are more strongly influenced by disorder. In previous studies, CD prevalence has often been assessed by estimating the percentage of a proteome containing a CD stretch greater than or equal to 30 amino acids in length [9, 14-16, 26, 27]. However, to our knowledge rigorously determined expected values and detailed ranges have not been reported numerically. Here, we assessed the distribution of the longest CD region (CD_L_) in each proteome.

The CD_L_ expected values varied from 42 amino acids (*A. thaliana*) to 104 amino acids (*D. melanogaster*), 26 amino acids (*C. reinhardtii*) to 41 amino acids (*D. melanogaster*), and 20 amino acids (*A. thaliana*) to 30 amino acids (*D. melanogaster*) for IUPred, DisEMBL-H, and DisEMBL-R predictors, respectively, with 20 of 30 expected values being greater than or equal to 30 amino acids (**Fig. 2**). Segment lengths of 8 amino acids (*A. thaliana*, IUPred), 16 amino acids (*C. reinhardtii*, DisEMBL - H), and seven amino acids (*D. discoideum*, DisEMBL - R) represented the lowest 25^th^ percentile values for the CD_L_, whereas the greatest 75^th^ percentile values were found to be 132 amino acids (*D. melanogaster*, IUPred), 46 amino acids (D. melanogaster, DisEMBL - H), and 40 amino acids (*D. melanogaster*, DisEMBL - R) (**Fig. 2**). Interestingly, the dispersion of the IUPred distributions was far greater than those of DisEMBL-H and DisEMBL-R. Lengths of 39 amino acids (*A. thaliana*) and 117 amino acids (*D. melanogaster*) were found to be the minimum and maximum IQR size for CD_L_ distributions predicted by IUPred (**Fig. 2A**), whereas the respective minimum-maximum IQR sizes for DisEMBL-H and DisEMBL-R were 15 (*C. reinhardtii*) to 26 amino acids (*D. melanogaster*) and 16 amino acids (*A. thaliana*) to 30 amino acids (*D. melanogaster*) (**Fig. 2B, C**). While the greatest 95^th^ percentile values for CD_L_ varied greatly from 50 amino acids (DisEMBL-R, *A. thaliana*) to 357 amino acids (IUPred, *D. melanogaster*), all (30 out of 30) were greater than or equal to 50 amino acids. Probability densities are shown in **Supplemental Fig. 3**. Explicit statistical values have been displayed in **Supplemental Table 4**.

**Fig 2.**
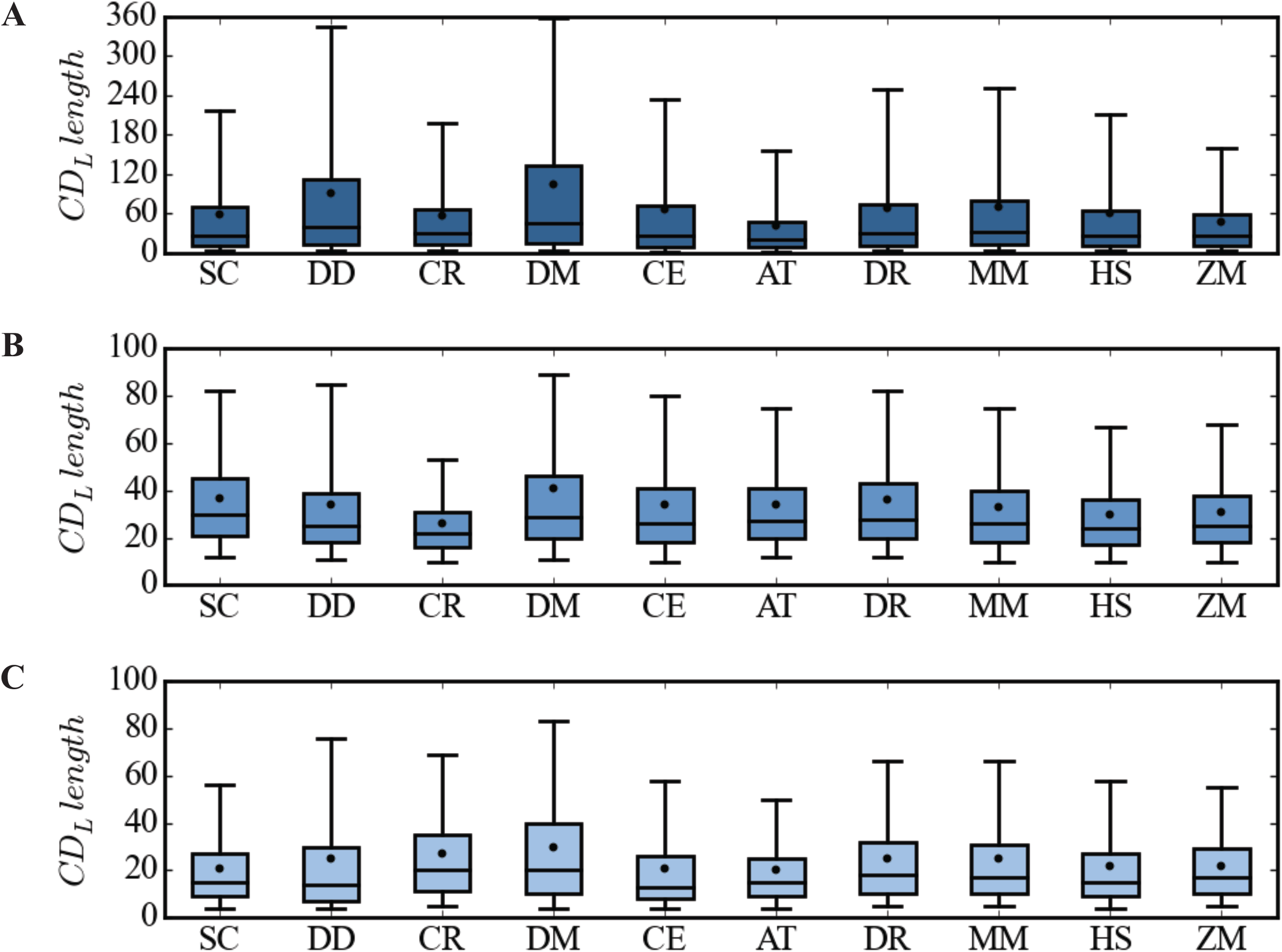
Longest CD stretch distribution in ten eukaryotic proteomes. Boxplots of CD_L_ regions calculated analyzing the predictions from IUPred (A), DisEMBL-H (B), and DisEMBL-R (C) are shown. The horizontal line within the box indicates the median, whereas the dots indicate the expected value determined via **Eq. 1**. The whiskers indicate the 5^th^ and 95^th^ percentile values. Numerical values are summarized in **Table 4**. Note that the minimum length CD segment is 2 amino acids, however, zero has been set as the minimum of the y-axis to preserve regularity for interpretation.

For the disorder prediction methods considered here, our results indicate that the significance threshold of 30 amino acids may only be appropriate when using DisEMBL-R, as this length often fails exceed the interquartile range for DisEMBL-H and IUPred predictions. Provided eight of the ten eukaryotes had a 75^th^ percentile for the CD_L_ region greater than or equal to 38 amino acids and 60 amino acids for DisEMBL-H and IUPred predictors, respectively, predictor-specific threshold values greater than the currently used 30 amino acids CD_L_ significance value should be established for these predictors (see **Discussion**). Furthermore, the differences in the magnitude of both the expected and 75^th^ percentile values observed here underscore the need to establish predictor-specific thresholds instead of adhering to a single, universal value.

### 3.4 In CD-containing proteins, the longest disordered region accounts for 6-18% of a protein’s total length

While the results presented in **Fig. 2** provide useful, intuitive information regarding the typical length expected of the CD_L_ region contained within a protein, it is subject to nebulous interpretations as it is not in the context of primary sequence length. To address this issue, we next analyzed the percentage of the total length of a protein that is accounted for by the longest continuously disordered segment. For each CD-containing protein, the longest CD percentage of length (LCPL) was simply calculated by dividing the length of the CD_L_ by the primary sequence length and multiplying the result by one hundred. LCPL distributions were analyzed in each proteome and expected values were obtained via **Eq. 1**.

For IUPred, DisEMBL – H, and DisEMBL – R, the expected values for LCPL varied from ∼12% (*A. thaliana*) to 18% (*H. sapiens*), ∼9% (*D. melanogaster*) to 16% (*H. sapiens*), and ∼6% (*S. cerevisiae*) to 10% (*H. sapiens*), respectively (**Fig. 3**). The minimum 25^th^ percentile values were found to be ∼2% (*A. thaliana*), ∼4% (*D. melanogaster*), and ∼2% (*D. discoideum*), with maximum 75^th^ percentile values of ∼24% (*H. sapiens*), ∼20% (*H. sapiens*), and ∼12% (*H. sapiens*) for IUPred, DisEMBL – H, and DisEMBL – R, respectively (**Fig. 3**). The organisms defining the boundaries of the aforementioned LCPL expected value ranges also exhibited the least (*H. sapiens*, IUPred; *D. melanogaster*, DisEMBL – H; *S. cerevisiae*, DisEMBL-R) and most (*H. sapiens*, all predictors) dispersion within the central 50% of the population (**Fig. 3**). Furthermore, we found the 95^th^ percentile for LCPL to vary dramatically between predictors, as it ranged from ∼17% (*S. cerevisiae*, DisEMBL-R) to ∼76% (*H. sapiens*, IUPred) (**Fig. 3**). Provided all of the expected values, and 25 out of 30 of the 75^th^ percentile values are below 20%, our results suggest the CD_L_ segment contained in a protein is typically less than 20% of the total protein length, and CD segments occupying a greater percentage of the total length may be significant. Probability densities can be found in **Supplemental Fig. 4**. Explicit statistical values have been displayed in **Supplemental Table 5**.

**Fig 3.**
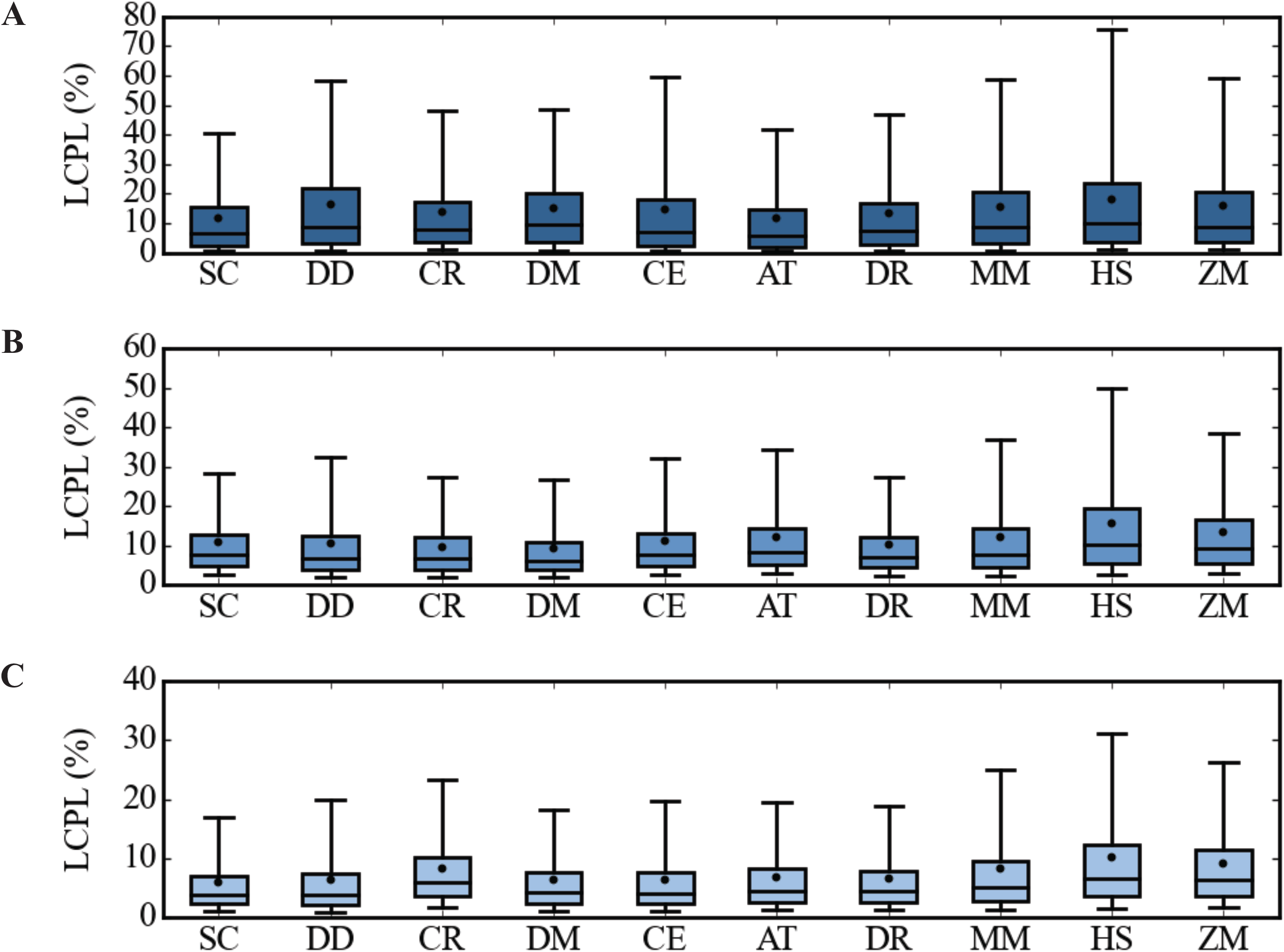
Longest CD percentage of length (LCPL) distribution in ten eukaryotic proteomes. Boxplots of LCPL determined by IUPred (A), DisEMBL-H (B), and DisEMBL-R (C) are shown. The horizontal line within the box indicates the median, whereas the dots indicate the expected value determined via **Eq. 1**. The whiskers represent the 5^th^ and 95^th^ percentiles. Numerical values are summarized in **Table 4**.

### 3.5 The reliability of significance thresholds for CD length varies between predictors in a protein length-dependent fashion

While the length of an intrinsically disordered region offers little insight into the purpose, function, or importance of that intrinsically disordered region, it has nevertheless become common practice to examine CD region length when using computational methods to identify, and/or assess the prevalence of, significantly long intrinsically disordered regions. When assessing the prevalence of significantly long CD regions, the percentage of CD segments greater than or equal to a fixed length is often examined, with 30 amino acids representing the commonly used value [9, 14-16, 26, 27]. However, when deeming a CD region as significant on the basis of length alone, utilizing a fixed threshold length becomes less reliable as the primary sequence length increases. For instance, although intrinsically disordered region length has not been found to specifically determine function in intrinsically disordered proteins, many would be more willing to accept that a CD region of 40 amino acids contributes more to the overall character of a protein that is 200 amino acids long compared to the same length segment in a protein with a primary sequence length of 2,000 amino acids, as this region accounts for a greater percentage of the length in the former protein and is arguably more likely to hold greater influence over structure (or lack-of-structure) overall. This consideration leads us to arrive at the following question. When is a protein too long to evaluate the significance of a CD region on the basis of its length alone?

To answer this question, we determined the protein length at which the CD_L_ region expected value (**Fig. 2**) begins to fall below the 25^th^ percentile cutoff for the LCPL (**Fig. 3**) (the concept of this threshold protein length is depicted in **Fig. 4A**). These values were obtained by solving **Eq. 2** for the protein length:

**Fig. 4.**
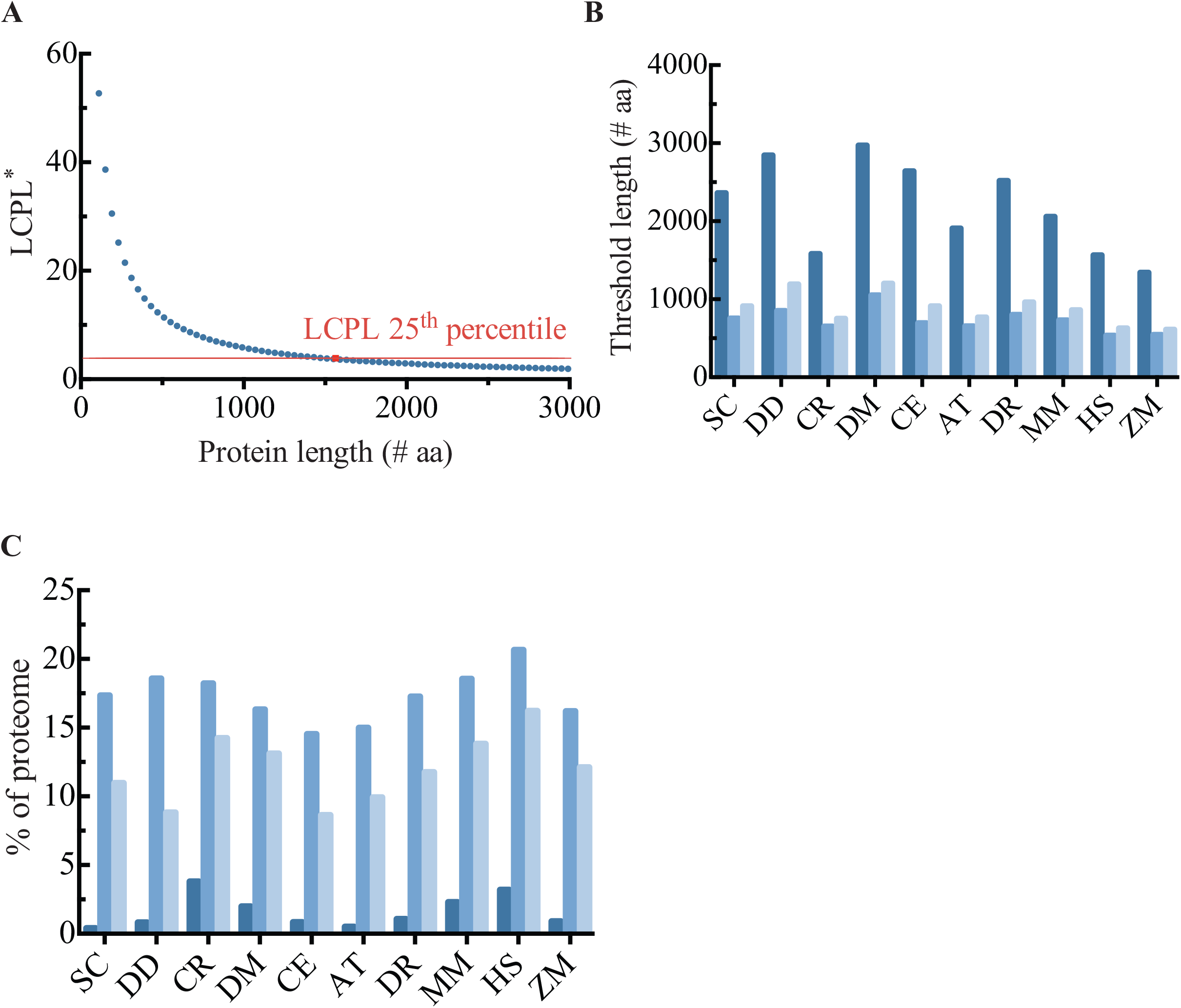
Reliability of CD length thresholds with increasing protein length. (A) Graphical depiction of the concept for estimating LCPL protein length thresholds. **Eq. 2** was solved to find the protein length where the LCPL begins to fall below the 25^th^ percentile LCPL value specific to the proteome and prediction algorithm. The red dot is the IUPred protein length threshold result in the *Homo sapiens* proteome. The dark blue dots are LCPL values calculated with Eq. 2 using protein lengths from 110 to 3,000 amino acids and are for conceptual purposes only. (B) Protein length threshold (PLT) values marking the maximum protein length where CD regions can be considered significant on the basis of length alone. Numeric values are provided in **Table 4**. (C) Percentage of the CD-containing proteins of each proteome with a length greater than or equal to the threshold values displayed in (B). Results for all proteomes are displayed for IUPred (dark blue), DisEMBL-H (medium blue), and DisEMBL-R (light blue).

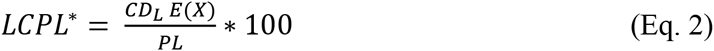

In **Eq. 2**, LCPL^*^ is the LCPL 25^th^ percentile value, CD_L_ E(X) is the expected value determined for the longest CD region, and PL is the primary sequence length. **Fig. 4A** illustrates the concept used for a single predictor in a single proteome, whereas the results for all predictors in the CD-containing proteins of all proteomes are displayed in **Fig. 4B**.

Protein length threshold (PLT) values ranged from 1,343-2,971 amino acids, 536-1,051 amino acids, and 611-1,200 amino acids for IUPred, DisEMBL-H, and DisEMBL-R, respectively (**Fig. 4B**). In all cases, PLT values were found to be substantially longer for IUPred compared to DisEMBL (**Fig. 4B**). For DisEMBL methods, the CD_L_ *E(X)* predicted by DisEMBL-R was found to be more tolerant of longer primary sequences than the DisEMBL-H algorithm (**Fig. 4B**). While the percentage of proteomes (specifically, the CD-containing population of each proteome) consisting of proteins with a primary sequence length greater than the predictor-specific PLT was low for IUPred (primarily due to higher CD_L_ E(X)), it was substantially greater for DisEMBL predictions, as it exceeded 10-15% in many cases (**Fig. 4C**). Thus, this result suggests that while the deficiency inherent to the CD_L_ metric may be of little concern when using IUPred, greater attention must be given to protein length when assessing CD significance using DisEMBL.

### 3.6 LCPL is stricter than CD_L_ when gauging the significance of continuous disordered regions in long proteins

We subsequently compared the effect of two different length thresholds (the commonly used 30 amino acids value and the CD_L_ 75^th^ percentile values (**Fig. 2**)), as well as the LCPL 75^th^ percentile values (**Fig. 3**), in identifying proteins with a CD region of significant length. Specifically, this analysis was performed in the subpopulation of CD containing proteins with a primary sequence length greater than or equal to the prediction algorithm-specific threshold primary sequence length determined for each proteome (**Fig. 4B**).

Between the two length thresholds, a smaller percentage of each subpopulation was predicted to contain a significantly long disordered segment when using the LCPL 75^th^ percentile determined in this study (**Fig. 5A, C**), although this was more variable with DisEMBL-R due to the more conservative nature of its CD predictions (**Fig. 5E**). Nevertheless, a substantial fraction of each subpopulation was still found to contain the feature of interest (**Fig. 5A, C, E**), supporting the **Fig. 4** assertion that conferring significance to a CD region on the basis of raw length alone is inappropriate for proteins with a residue count exceeding the algorithm-specific PLT. We subsequently explored the effect of classification using the LCPL metric as guidance. From **Fig. 5B, D**, and **F**, a substantial decrease in the percentage of proteins exhibiting a significant CD_L_ segment is observed when using the LCPL for classification. In all cases, less than 15% of the PLT-exceeding subpopulation can be considered as having a significantly long CD region, whereas the same was found for less than 10% of the DisEMBL subpopulations (**Fig. 5B, D, F**). Taken together, these results exemplify the point that the LCPL is an excellent metric to guide scientists evaluating significantly long disordered regions in proteins having lengths exceeding the predictor-specific threshold values determined in **Fig. 4**.

**Fig. 5.**
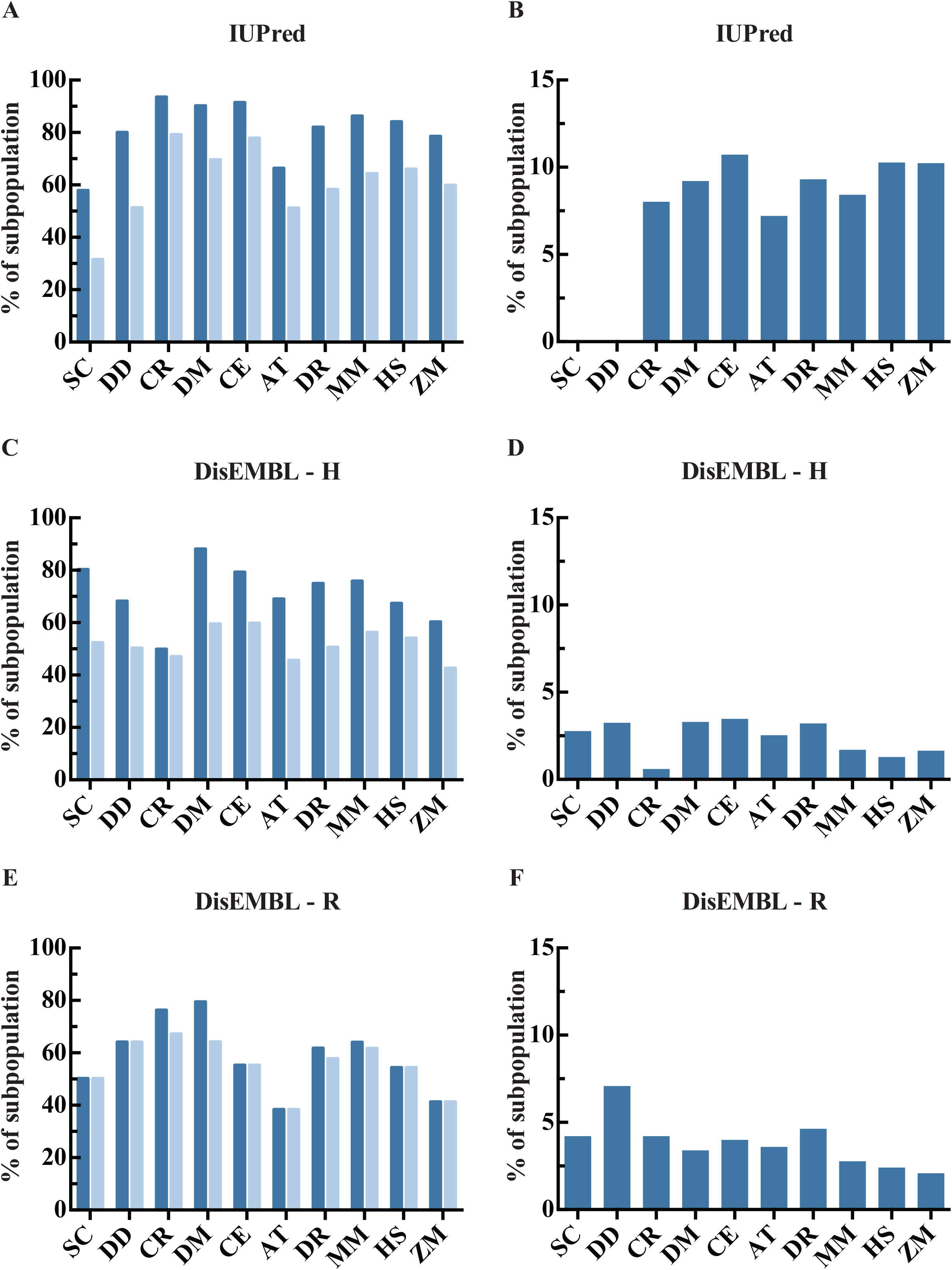
LCPL is a strict metric for gauging the significance of continuous disordered regions in long proteins. The subpopulation of proteins having a length greater than or equal to the predictor-specific protein length threshold presented in **Fig. 4** was analyzed. (A, C, E) The percentage of the subpopulation containing a longest continuous disordered region greater than or equal to 30 amino acids (dark blue bars), or the proteome-specific 75^th^ percentile value (light blue bars). (B, D, F) The percentage of the subpopulation having a LCPL value greater than or equal to the proteome-specific 75^th^ percentile value. Results are displayed for IUPred (A, B), DisEMBL-H (C, D), and DisEMBL-R (E, F).

### 3.7 Example usage of the presented guidelines

To illustrate how these guidelines might be utilized, we will briefly assess disorder using DisEMBL - H in three different proteins taken from the *Homo sapiens* proteome: α-synuclein (UniProt AC: P37840), MARCH7 (UniProt AC: Q9H992), and the Protein transport protein Sec61 subunit α isoform 1 (UniProt AC: P61619). α-synuclein is an intrinsically disordered presynaptic neuronal protein [28]. Using our guidelines, α-synuclein is indeed classified as highly disordered with its disorder content in the 89^th^ percentile (**Fig. 6 A, D**). Furthermore, even though the CD_L_ of 28 amino acids in α-synuclein is not significant with respect to the rest of the proteome (**Fig. 6 B, D**), this region does account for a significant fraction of the protein (**Fig. 6 C, D**), which could serve to hint that this region has a substantial influence over the lack of structure observed in this protein. Next we analyze MARCH7, an E3 ubiquitin-protein ligase with a primary sequence length of 704 amino acids, which exceeds the DisEMBL – H PLT of 536 amino acids determined for the Homo sapiens proteome (**Table 4**). The disorder content and CD_L_ of MARCH7 were respectively found to be in the 85^th^ and 93rd percentiles (**Fig. 6 A, B, D**), although the CD_L_ was not found to account for a substantial fraction of MARCH7 (Fig. 6 C, D), making it difficult to interpret the importance of this region overall. Lastly, Protein transport protein Sec61 subunit α isoform 1 is a 476 amino acid long protein, known to be a highly conserved component of the endoplasmic reticulum translocon complex that serves as the high traffic [29] gateway into the ER [30-32]. Employing our guidelines, Protein transport protein Sec61 subunit α isoform 1 is predicted by DisEMBL – H to be ordered, with percent disorder and LCPL below the 25^th^ percentile value (**Fig. 6 A, C, D**), and its CD_L_ in the 40^th^ percentile (**Fig. 6 B, D**).

**Fig. 6.**
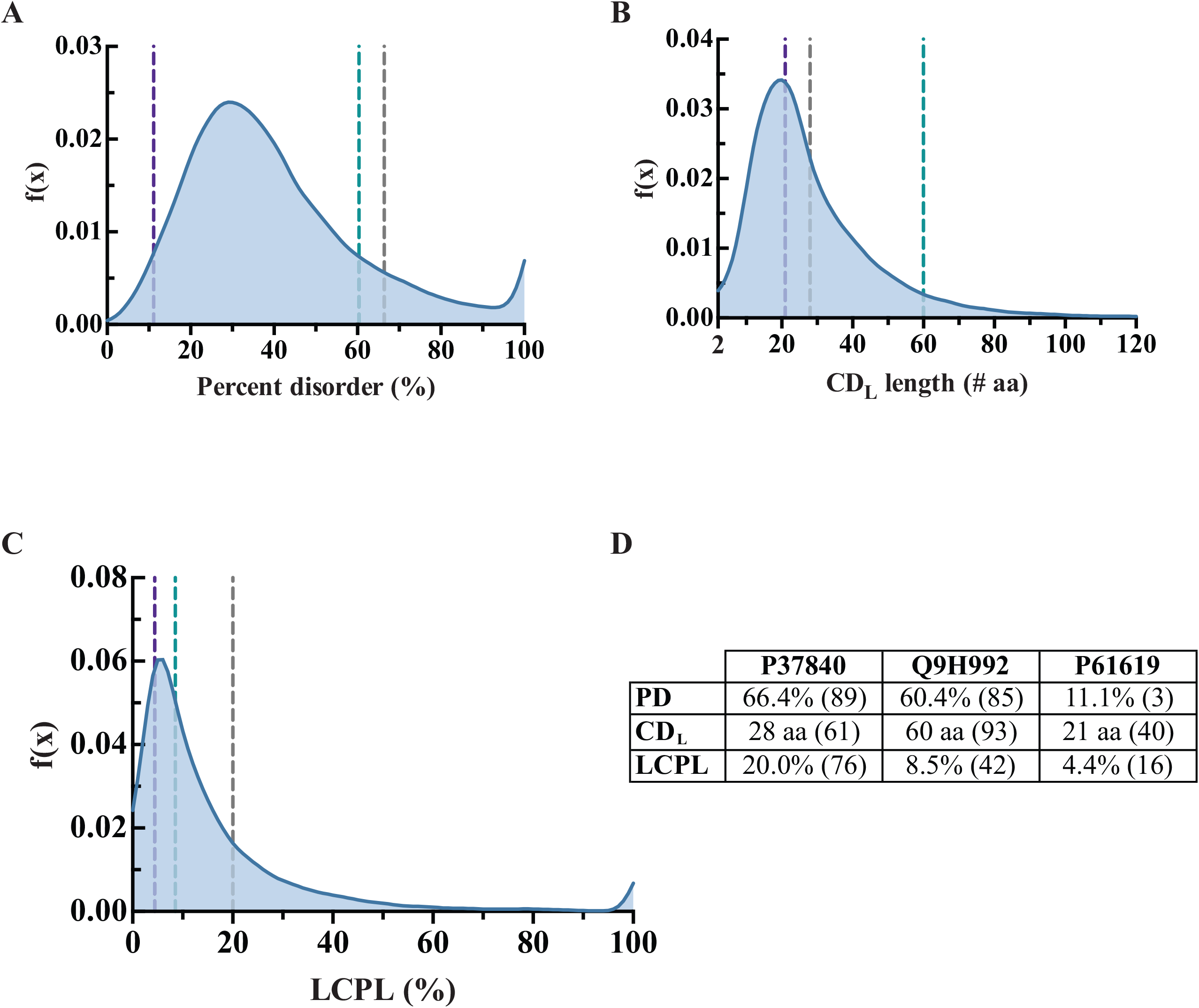
Example of a standardized assessment of disorder based on our proteome-based descriptive statistics guidelines. Disorder content (A), CD_L_ (B), and LCPL (C) were assessed with DisEMBL – H in α-synuclein (P37840, gray vertical line), MARCH7 (Q9H992, teal vertical line), and the Protein transport protein Sec61 subunit α isoform 1 (P61619, purple vertical line). The values have been displayed in (D), with the percentile values in parentheses. For (B), the truncated range of [2, 120] has been displayed to facilitate visualization, as the density beyond 120 amino acids was near zero.

**Table 4.**
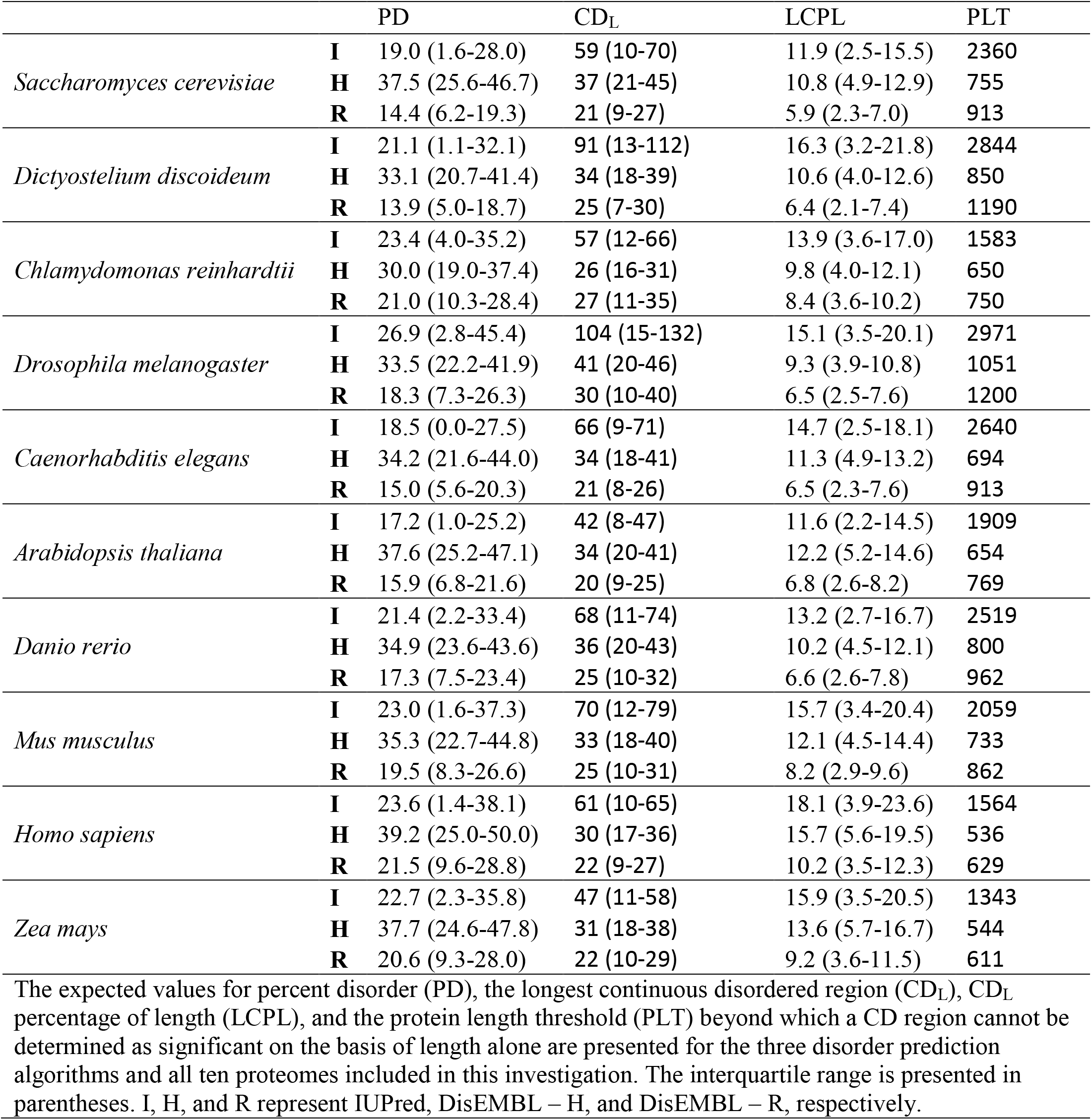
Summary of proteome-based descriptive guidelines for disorder characterization.

|  |  | PD | CD<sub>L</sub> | LCPL | PLT |
| --- | --- | --- | --- | --- | --- |
| *Saccharomyces cerevisiae* | **I** | 19.0 (1.6-28.0) | 59 (10-70) | 11.9 (2.5-15.5) | 2360 |
|  | **H** | 37.5 (25.6-46.7) | 37 (21-45) | 10.8 (4.9-12.9) | 755 |
|  | **R** | 14.4 (6.2-19.3) | 21 (9-27) | 5.9 (2.3-7.0) | 913 |
| *Dictyostelium discoideum* | **I** | 21.1 (1.1-32.1) | 91 (13-112) | 16.3 (3.2-21.8) | 2844 |
|  | **H** | 33.1 (20.7-41.4) | 34 (18-39) | 10.6 (4.0-12.6) | 850 |
|  | **R** | 13.9 (5.0-18.7) | 25 (7-30) | 6.4 (2.1-7.4) | 1190 |
| *Chlamydomonas reinhardtii* | **I** | 23.4 (4.0-35.2) | 57 (12-66) | 13.9 (3.6-17.0) | 1583 |
|  | **H** | 30.0 (19.0-37.4) | 26 (16-31) | 9.8 (4.0-12.1) | 650 |
|  | **R** | 21.0 (10.3-28.4) | 27 (11-35) | 8.4 (3.6-10.2) | 750 |
| *Drosophila melanogaster* | **I** | 26.9 (2.8-45.4) | 104 (15-132) | 15.1 (3.5-20.1) | 2971 |
|  | **H** | 33.5 (22.2-41.9) | 41 (20-46) | 9.3 (3.9-10.8) | 1051 |
|  | **R** | 18.3 (7.3-26.3) | 30 (10-40) | 6.5 (2.5-7.6) | 1200 |
| *Caenorhabditis elegans* | **I** | 18.5 (0.0-27.5) | 66 (9-71) | 14.7 (2.5-18.1) | 2640 |
|  | **H** | 34.2 (21.6-44.0) | 34 (18-41) | 11.3 (4.9-13.2) | 694 |
|  | **R** | 15.0 (5.6-20.3) | 21 (8-26) | 6.5 (2.3-7.6) | 913 |
| *Arabidopsis thaliana* | **I** | 17.2 (1.0-25.2) | 42 (8-47) | 11.6 (2.2-14.5) | 1909 |
|  | **H** | 37.6 (25.2-47.1) | 34 (20-41) | 12.2 (5.2-14.6) | 654 |
|  | **R** | 15.9 (6.8-21.6) | 20 (9-25) | 6.8 (2.6-8.2) | 769 |
| *Danio rerio* | **I** | 21.4 (2.2-33.4) | 68 (11-74) | 13.2 (2.7-16.7) | 2519 |
|  | **H** | 34.9 (23.6-43.6) | 36 (20-43) | 10.2 (4.5-12.1) | 800 |
|  | **R** | 17.3 (7.5-23.4) | 25 (10-32) | 6.6 (2.6-7.8) | 962 |
| *Mus musculus* | **I** | 23.0 (1.6-37.3) | 70 (12-79) | 15.7 (3.4-20.4) | 2059 |
|  | **H** | 35.3 (22.7-44.8) | 33 (18-40) | 12.1 (4.5-14.4) | 733 |
|  | **R** | 19.5 (8.3-26.6) | 25 (10-31) | 8.2 (2.9-9.6) | 862 |
| *Homo sapiens* | **I** | 23.6 (1.4-38.1) | 61 (10-65) | 18.1 (3.9-23.6) | 1564 |
|  | **H** | 39.2 (25.0-50.0) | 30 (17-36) | 15.7 (5.6-19.5) | 536 |
|  | **R** | 21.5 (9.6-28.8) | 22 (9-27) | 10.2 (3.5-12.3) | 629 |
| *Zea mays* | **I** | 22.7 (2.3-35.8) | 47 (11-58) | 15.9 (3.5-20.5) | 1343 |
|  | **H** | 37.7 (24.6-47.8) | 31 (18-38) | 13.6 (5.7-16.7) | 544 |
|  | **R** | 20.6 (9.3-28.0) | 22 (10-29) | 9.2 (3.6-11.5) | 611 |

The expected values for percent disorder (PD), the longest continuous disordered region (CD<sub>L</sub>), CD<sub>L</sub> percentage of length (LCPL), and the protein length threshold (PLT) beyond which a CD region cannot be determined as significant on the basis of length alone are presented for the three disorder prediction algorithms and all ten proteomes included in this investigation. The interquartile range is presented in parentheses. I, H, and R represent IUPred, DisEMBL-H, and DisEMBL-R, respectively.

## 4. Discussion

We presented a thorough analysis of intrinsic disorder predicted by two reputable algorithms based on physicochemical principles. Our analysis utilized a non-parametric statistical approach to estimate predictor-specific objective guidelines for determining whether the disorder content or length of a CD region is significant with respect to the rest of the proteomic population. A summary of our guidelines is displayed in **Table 4**. The complete statistical assessment for percent disorder, CD_L_, and LCPL can be found in **Supplemental Tables 3-5**.

Expected values and ranges were found to vary between the different disorder prediction methods. DisEMBL-R, an artificial neural network trained on missing electron density assignments in the Protein Data Bank, consistently predicted fewer disordered residues than did IUPred or DisEMBL-H (**Fig. 1**). Overall, the disorder content ranges for IUPred and DisEMBL-R were in agreement with a range of ∼16-22% [14] and an average of 20.5% [16] reported by two proteomic investigations of intrinsic disorder, whereas DisEMBL-H disorder content predictions were found to be substantially higher, but still consistent with results from a third proteomic disorder investigation [15](**Fig. 1**). Moreover, although DisEMBL-H predicted more disorder than DisEMBL-R, it does so with greater accuracy given its lower false positive rate [24], as disorder only represents one potential cause for missing coordinates in the X-ray crystallography data used to train DisEMBL-R. Regarding the IUPred predictions, it is important to understand that the low 25th percentile values observed in the IUPred disorder content distributions (**Fig. 1A**) prevents the use of its 25^th^ percentile as a mark above which proteins begin to lack significant order. Regardless, the 25^th^ percentile values are interpretable for DisEMBL predictions, and in all cases, the 75^th^ percentile values can be used to define proteins containing significantly high disorder content.

To provide insight into the organization of disordered residues, the distributions of the longest CD region (CD_L_) and the percentage of residues accounted for by the longest continuous disordered region (LCPL) were examined. Due to the restraints imposed by the minimum theoretical value of a CD region (2 amino acids), we limited all CD analyses to the CD-containing population of each proteome (**Table 3**), resulting in the exclusion of completely ordered proteins, as well as proteins exhibiting disorder exclusively in the form of isolated amino acids. While this may lead to inflation of the disorder features with respect to the whole proteome population, a large fraction of each proteome (≥70% in all cases) was found to exhibit CD, as predicted by each algorithm (**Table 3**). Therefore we believe that our sample protein populations are representative, and dismiss any inflation resulting from the enforcement of the eligibility criterion to be minimal. Additionally, we acknowledge that the CD_L_ metric does not account for various features contributing to the overall disorder content of a protein, such as isolated disordered amino acids and/or shorter CD segments that may exist in areas nearby the CD_L_ region or other CD regions. Regardless, CD_L_ provides an indication of the most organized disordered segment in a protein, and when examined at the population level, it offers valuable insight into the largest disordered segment one should expect in a given protein. IUPred predictions provided the greatest expected values with the most dispersion (**Fig. 2A**) when examining CD_L_ regions; this trend was also found in the LCPL distributions (**Fig. 3A**).

When deeming CD regions significant on the basis of length alone, it has become common practice to make this evaluation with respect to a universal fixed length. Numerous studies utilizing various disorder prediction algorithms have most commonly used a threshold value of 30 consecutive disordered residues to define a ‘long disordered region’ [9, 14-16, 27, 28]. For DisEMBL-R, this value appears to be a reasonable cutoff for significance, as none of the proteomes were found to have expected values greater than 30 amino acids and six of the ten proteomes were found to have 75^th^ percentile cutoffs less than or equal to 30 amino acids with the remaining four proteomes having 75^th^ percentile cutoffs less than or equal to 35 amino acids (**Table 4**). However, the threshold of 30 amino acids appears to be insufficient for identifying significantly long CD_L_ regions when using the IUPred and DisEMBL-H algorithms. This was suggested by the observation that all proteomes were found to have CD_L_ 75^th^ percentile values greater than 30 amino acids when assessing CD_L_ regions with IUPred and DisEMBL-H. Furthermore, all and nine of ten proteomes were found to contain expected values for CD_L_ length greater than 30 amino acids for IUPred and DisEMBL-H, respectively.

Considering the above, we suggested algorithm-specific thresholds be established that extend beyond the current 30 amino acids value, when identifying “significantly long” CD regions in a population on the basis of length alone. For DisEMBL-H, we propose that a CD_L_ threshold length of 40 amino acids would be more appropriate on the basis that eight of the ten eukaryotes exhibited 75^th^ percentile values greater or equal than 38 amino acids, with six of ten exceeding 40 amino acids. For IUPred, the threshold value should be set even higher, at 60 amino acids, as eight of the ten proteomes had 75^th^ percentile values greater than 60 amino acids (albeit the range of the IUPred CD_L_ 75^th^ percentile values was much greater than that of DisEMBL-H) (**Table 4**). The proposed increases would be substantial, with the DisEMBL-H and IUPred thresholds representing a 33% and 100% increase over the existing value of 30 amino acids.

One major concern of the CD_L_ guidance metric is that its power diminishes with increasing primary sequence length. To provide a means for identifying when a protein is too long for compatibility with this metric, we estimated protein length threshold values (referred to as a “PLT”), specific to each disorder prediction algorithm, and for all ten eukaryotes analyzed (**Fig. 4**). The prevalence of this issue was dramatically lower for IUPred predictions, given less than 5% of every proteome had a length greater than the predictor-specific PLT; whereas the issue was much more pronounced for DisEMBL predictions as over 10% of most proteomes were found to exceed the PLT length (**Fig. 4C**). When determining a conspicuous CD region in a protein exceeding these aforementioned threshold protein lengths, it is recommended that the LCPL metric be used in place of a length threshold, as we have shown the LCPL to be far more selective in general (**Fig. 5**).

Finally, we demonstrated the value of our guidelines in assessing disorder in three different proteins (**Fig. 6**). From the simple example presented, it can be observed that our guidelines, which are disorder prediction algorithm-specific and proteome-specific, provide a valuable tool for interpreting the significance of the output of the disorder prediction methods considered here.

## 5. Conclusions

The guidelines presented here are intended to facilitate biochemical and biophysical scientists in making simple objective disorder classifications in a protein of interest belonging to one of the ten eukaryotic proteomes included in our analysis (in addition to the **Table 3** summary, explicit descriptive statistics have been included in **Supplemental Tables 3-5**). Specifically, these guidelines are to be used with the IUPred and DisEMBL algorithms, and we further intend our analysis to inspire the creation of similar guidelines for alternative disorder prediction algorithms, as it is very difficult to assess the significance of disorder predictions without guidelines of this nature. We acknowledge that our analysis is descriptive; it aims to place disorder predictions into the context of the proteome and facilitate the identification of disorder anomalies when conducting an analysis of disorder in a protein of interest.

Not only can the guidelines reported here be used to evaluate the significance of disordered properties in an individual protein, but these guidelines can also greatly facilitate exploratory computational screens that seek to identify proteins with significant disordered features for further experimental investigation. Furthermore, although our study was limited to a small number of prediction tools, the general analytical approach is amenable to any disorder prediction algorithm with computational performance suitable for whole proteome analysis. Thus, a bigger picture goal of this work is that it will inspire similar analyses to be performed prior to the release of new disorder prediction algorithms, as well as for other existing algorithms, in order to facilitate the interpretation of disorder predictions. With a universal disorder prediction tool currently absent, together with the variation in disorder predictions observed between different algorithms and between different proteomes, the meaningful interpretation of disorder predictions relies heavily on guidelines like the ones presented in this work.

## Acknowledgements

We thank Dr. Kerby Shedden (Center for Statistical Consultation and Research, University of Michigan) for his technical support. We also thank Suzanne K. Shoffner for critically reading the manuscript. This work was partially supported by the University of Michigan Protein Folding Diseases Initiative and the University of Michigan Medical School Research Discovery Fund.

## References

[1] R.W. Kriwacki, L. Hengst, L. Tennant, S.I. Reed, P.E. Wright, Structural studies of p21Waf1/Cip1/Sdi1 in the free and Cdk2-bound state: conformational disorder mediates binding diversity, Proc Natl Acad Sci U S A, 93 (1996) 11504–11509.

[2] K.W. Plaxco, M. Grob, The importance of being unfolded, Nature, 386 (1997) 657–659.

[3] P.E. Wright, H.J. Dyson, Intrinsically unstructured proteins: re-assessing the protein structure-function paradigm, J Mol Biol, 293 (1999) 321–331.

[4] H.J. Dyson, P.E. Wright, Intrinsically unstructured proteins and their functions, Nat Rev Mol Cell Biol, 6 (2005) 197–208.

[5] P. Tompa, Intrinsically disordered proteins: a 10-year recap, Trends Biochem Sci, 37 (2012) 509–516.

[6] L.M. Iakoucheva, C.J. Brown, J.D. Lawson, Z. Obradović, A.K. Dunker, Intrinsic Disorder in Cell-signaling and Cancer-associated Proteins, Journal of Molecular Biology, 323 (2002) 573–584.

[7] P.E. Wright, H.J. Dyson, Intrinsically disordered proteins in cellular signalling and regulation, Nat Rev Mol Cell Biol, 16 (2015) 18–29.

[8] G.J. Rautureau, C.L. Day, M.G. Hinds, Intrinsically disordered proteins in bcl-2 regulated apoptosis, Int J Mol Sci, 11 (2010) 1808–1824.

[9] Z. Peng, B. Xue, L. Kurgan, V.N. Uversky, Resilience of death: intrinsic disorder in proteins involved in the programmed cell death, Cell Death Differ, 20 (2013) 1257–1267.

[10] M.K. Yoon, D.M. Mitrea, L. Ou, R.W. Kriwacki, Cell cycle regulation by the intrinsically disordered proteins p21 and p27, Biochem Soc Trans, 40 (2012) 981–988.

[11] V. Vittal, L. Shi, D.M. Wenzel, K.M. Scaglione, E.D. Duncan, V. Basrur, K.S. Elenitoba-Johnson, D. Baker, H.L. Paulson, P.S. Brzovic, R.E. Klevit, Intrinsic disorder drives N-terminal ubiquitination by Ube2w, Nat Chem Biol, 11 (2015) 83–89.

[12] B. Wang, S.A. Merillat, M. Vincent, A.K. Huber, V. Basrur, D. Mangelberger, L. Zeng, K. Elenitoba-Johnson, R.A. Miller, D.N. Irani, A.A. Dlugosz, S. Schnell, K.M. Scaglione, H.L. Paulson, Loss of the Ubiquitin-conjugating Enzyme Ube2W results in susceptibility to early postnatal lethality and defects in skin, immune and male reproductive systems, J Biol Chem, 291 (2015) 3030–3042.

[13] T.P. Knowles, M. Vendruscolo, C.M. Dobson, The amyloid state and its association with protein misfolding diseases, Nat Rev Mol Cell Biol, 15 (2014) 384–396.

[14] J.J. Ward, J.S. Sodhi, L.J. McGuffin, B.F. Buxton, D.T. Jones, Prediction and functional analysis of native disorder in proteins from the three kingdoms of life, J Mol Biol, 337 (2004) 635–645.

[15] B. Xue, A.K. Dunker, V.N. Uversky, Orderly order in protein intrinsic disorder distribution: disorder in 3500 proteomes from viruses and the three domains of life, J Biomol Struct Dyn, 30 (2012) 137–149.

[16] Z. Peng, J. Yan, X. Fan, M.J. Mizianty, B. Xue, K. Wang, G. Hu, V.N. Uversky, L. Kurgan, Exceptionally abundant exceptions: comprehensive characterization of intrinsic disorder in all domains of life, Cell Mol Life Sci, 72 (2015) 137–151.

[17] C.L. Ogden, M.D. Carroll, B.K. Kit, K.M. Flegal, Prevalence of obesity and trends in body mass index among US children and adolescents, 1999-2010, JAMA, 307 (2012) 483–490.

[18] K.M. Flegal, M.D. Carroll, B.K. Kit, C.L. Ogden, Prevalence of obesity and trends in the distribution of body mass index among US adults, 1999-2010, JAMA, 307 (2012) 491–497.

[19] J.S. Schiller, J.W. Lucas, B.W. Ward, J.A. Peregoy, Summary health statistics for U.S. adults: National Health Interview Survey, 2010, Vital Health Stat 10, (2012) 1–207.

[20] C. UniProt, UniProt: a hub for protein information, Nucleic Acids Res, 43 (2015) D204–212.

[21] B.E. Suzek, H. Huang, P. McGarvey, R. Mazumder, C.H. Wu, UniRef: comprehensive and non-redundant UniProt reference clusters, Bioinformatics, 23 (2007) 1282–1288.

[22] Z. Dosztanyi, V. Csizmok, P. Tompa, I. Simon, The pairwise energy content estimated from amino acid composition discriminates between folded and intrinsically unstructured proteins, J Mol Biol, 347 (2005) 827–839.

[23] Z. Dosztanyi, V. Csizmok, P. Tompa, I. Simon, IUPred: web server for the prediction of intrinsically unstructured regions of proteins based on estimated energy content, Bioinformatics, 21 (2005) 3433–3434.

[24] R. Linding, L.J. Jensen, F. Diella, P. Bork, T.J. Gibson, R.B. Russell, Protein disorder prediction: implications for structural proteomics, Structure, 11 (2003) 1453–1459.

[26] M.C. Jones, Simple boundary correction for kernel density estimation, Statistics and Computing, 3 (1993) 135–146.

[26] C. Haynes, C.J. Oldfield, F. Ji, N. Klitgord, M.E. Cusick, P. Radivojac, V.N. Uversky, M. Vidal, L.M. Iakoucheva, Intrinsic disorder is a common feature of hub proteins from four eukaryotic interactomes, PLoS Comput Biol, 2 (2006) e100.

[27] Z. Peng, M.J. Mizianty, L. Kurgan, Genome-scale prediction of proteins with long intrinsically disordered regions, Proteins, 82 (2014) 145–158.

[28] P.H. Weinreb, W. Zhen, A.W. Poon, K.A. Conway, P.T. Lansbury, Jr., NACP, a protein implicated in Alzheimer’s disease and learning, is natively unfolded, Biochemistry, 35 (1996) 13709–13715.

[29] M. Vincent, M. Whidden, S. Schnell, Surveying the floodgates: estimating protein flux into the endoplasmic reticulum lumen in Saccharomyces cerevisiae, Front Physiol, 5 (2014) 444.

[30] R.J. Deshaies, R. Schekman, A yeast mutant defective at an early stage in import of secretory protein precursors into the endoplasmic reticulum, J Cell Biol, 105 (1987) 633–645.

[33] R.J. Deshaies, R. Schekman, SEC62 encodes a putative membrane protein required for protein translocation into the yeast endoplasmic reticulum, J Cell Biol, 109 (1989) 2653–2664.

[32] E. Hartmann, T. Sommer, S. Prehn, D. Gorlich, S. Jentsch, T.A. Rapoport, Evolutionary conservation of components of the protein translocation complex, Nature, 367 (1994) 654–657.

